# A comparison of the parasitoid wasp species richness of tropical forest sites in Peru and Uganda – subfamily Rhyssinae (Hymenoptera: Ichneumonidae)

**DOI:** 10.1101/2023.08.23.554460

**Authors:** Hopkins Tapani, Tuomisto Hanna, Isrrael C. Gómez, Ilari E. Sääksjärvi

**Affiliations:** Zoological Museum, Biodiversity Unit, FI-20014 University of Turku, Finland; Department of Biology, FI-20014 University of Turku, Finland; Department of Biology, Aarhus University, Denmark

**Keywords:** biodiversity, idiobiont parasitoids, latitudinal diversity gradient, Malaise trap, Uganda Malaise trapping 2014-2015, Amazon Malaise trapping 2000

## Abstract

The global distribution of parasitoid wasp species richness is poorly known. Past attempts to compare data from different sites have been hampered by small sample sizes and lack of standardisation. During the past decades, we have carried out long-term Malaise trapping using a standardised approach in the tropical forests of Peru (western Amazonia) and Uganda (eastern Africa). Here, we test how well such data can be used for global comparisons, by comparing the results for the subfamily Rhyssinae (Hymenoptera: Ichneumonidae). We found that more rhyssine species were caught in Peru than in Uganda, despite the Ugandan samples containing many more individuals both in absolute terms and per unit time. The difference in the number of individuals caught may largely be due to more rainfall in Peru, since rain reduces Malaise trap catches. Peruvian traps caught species at a faster rate (per individual caught) than Ugandan traps. We interpret this as a sign that the Peruvian sites have more species than the Ugandan site. Long-term, standardised Malaise trapping showed promise for global comparisons of species richness. Sampling more sites on both continents, and analysing all subfamilies, would give an estimate of which continent has more parasitoid wasp species. We suggest some refinements to the sampling design that would further improve sampling efficiency for future studies.

## Introduction

Darwin wasps or ichneumonids (parasitoid wasps of the family Ichneumonidae) were once believed to display an “anomalous latitudinal diversity gradient”: to be more species rich in mid and high latitudes than in the tropics (Owen & Owen, 1974; Janzen & Pond, 1975; Janzen, 1981), instead of peaking in the tropics as is the case with most taxa (Hawkins, 2001; Willig et al., 2003). Although this belief has lost ground after the discovery of numerous unknown tropical Darwin wasp species, especially in Central America and South America (Gauld, 1991; Gaston & Gauld, 1993; Sääksjärvi et al., 2004), we still do not have enough data to draw reliable conclusions on where the species richness of Darwin wasps peaks (Quicke, 2012). We especially do not know how other tropical areas compare to the unexpectedly species-rich South American sites.

Long-term Malaise trapping is one potential way of finding out how the species richness of flying insects is distributed on our planet. Malaise traps are tent-like passive traps that collect flying insects, especially Diptera and Hymenoptera, giving large sample sizes with relatively little effort in terms of person-hours (van Achterberg, 2009; Saunders & Ward, 2018). Since Malaise traps are widely used and are available commercially in standard sizes and colourations, the potential for getting comparable samples from different sites is high. However, large sample sizes and several traps may be needed since the catches tend to vary greatly. Malaise traps typically catch quite different numbers of individuals even when placed near each other in the same habitat, depending on how their position relates to popular insect flight routes (Fraser et al., 2008; Saunders & Ward, 2018; Chimeno et al. 2023). Weather also affects catches: traps typically catch less in rainy weather, due to flying insects being less active, which can be hard to disentangle from genuine, seasonal changes in abundance (e.g. Hopkins, Roininen, & Sääksjärvi, 2019b). However, it is possible to estimate by mathematical modelling how large a catch (number of individuals of each species) to expect in given weather conditions and habitat. Such modelling has been done for the Ugandan site of the present paper (modelling described in Hopkins, Roininen, & Sääksjärvi, 2019b).

Despite the potential of long-term Malaise trapping for faunistic comparisons, few attempts have been made to compare the Malaise trapped Darwin wasp faunas of different sites. Timms et al. (2016) took 38 Darwin wasp datasets which had been gathered by Malaise trapping between latitudes 82°N and 25°S. Although they got some interesting results indicating that the latitudinal pattern displayed by Darwin wasps varies between subfamilies, only four of the datasets had been identified to species level, and those sites were so undersampled that the number of species and the number of individuals caught were interchangeable (figure 3 in their paper, although they still interpreted that species richness can be inferred from the observed abundances). Also, it is worth noting that the number of traps used at different sites was not taken into account: the apparently higher abundance in the tropics (e.g. figure 4a in the paper), for example, simply reflects the fact that more traps were used in the tropics than at higher latitudes. Gómez et al. (2017) compared the data of 97 sites on three different continents. Although their results tentatively suggested that the species richness of the subfamilies Pimplinae and Rhyssinae might peak in the tropics, the sample sizes were too small for firm conclusions. There was also a lot of variation in how the Malaise trapping had been conducted: for example, the Ugandan Kibale site in their data had been sampled with unusually small Malaise traps (Hopkins et al., 2018). The general picture emerging from these two attempts is that comparing the Darwin wasp species richness of different sites is challenging, due to most Malaise trapping having yielded small sample sizes and not having been conducted in a standardised way.

Recently, we have published the first results of long-term Malaise trapping in Kibale National Park, Uganda (Hopkins, Roininen, van Noort, et al., 2019; Hopkins, Roininen, & Sääksjärvi, 2019b). These results are on the relatively rarely caught rhyssine wasps (subfamily Rhyssinae), the only subfamily in the material that is currently sorted, pinned and otherwise ready to be studied. Since the methodology is the same as in our earlier long-term sampling in Peru (Sääksjärvi et al., 2004; Gómez et al., 2015), this allows us to compare the species richness of tropical forest sites on two continents, with material that has reasonable sample sizes and has been collected in a standardised way. The subfamily Rhyssinae is cosmopolitan in distribution and moderately small in terms of species richness. Rhyssines are idiobiont ectoparasitoids of holometabolous insects. They are mostly large in size and vividly coloured, and the females possess a long ovipositor for ovipositing into hosts living deeply concealed in decaying wood (e.g. Siricoidea wasps and wood-boring beetles).

In this work, we compare the abundance and species richness of rhyssine wasps in Ugandan and Peruvian Malaise trap samples. Our aim is to explore if similar analysis of other (more abundant) subfamilies, and further sampling at more sites, would allow the species richness of African and South American tropical forests to be compared.

## Methods

### Study sites

The two main study sites were Allpahuayo–Mishana National Reserve in north-eastern Peru (western Amazonia, South America), and Kibale National Park in Uganda (eastern Africa). Both are near the equator and predominantly covered by tropical forest. We also included data from a second western Amazonian site, Los Amigos Conservation Concession in south-eastern Peru.

Allpahuayo–Mishana National Reserve contains moist tropical forest that is known for its habitat heterogeneity (Whitney & Alonso, 1998; Sääksjärvi et al., 2004). The reserve is about 25 km southwest from the city of Iquitos (3°57 S, 73°26 W, approx. 110–180 m.a.s.l.: Gómez et al., 2015). We broadly classified the non-inundated forest (*tierra firme*) into forest types based on soil characteristics (Sääksjärvi et al., 2006; Hopkins, Gómez, et al., 2023). Mean annual rainfall is approximately 3000 mm and mean annual temperature is 26°C (Sääksjärvi et al., 2006). Weather data, consisting of daily rainfall and daily mean temperatures, were available for Iquitos city from NOAA/NCEI (https://www.ncdc.noaa.gov, Menne et al., 2012). In the analyses, we replaced any missing rain data with the 29-day average rainfall (average over the time period from 14 days before to 14 days after the day whose rain datum was missing). The study site is described in greater detail in our earlier papers (Sääksjärvi et al., 2004, 2006; Gómez et al., 2015) and one of the datasets associated with the present paper (Hopkins, Gómez, et al., 2023).

Los Amigos Conservation Concession contains moist tropical forest growing on a mosaic of different soils (Gómez et al., 2017). Our study site was near the Los Amigos Biological Station (CICRA, 12°34 S, 70°05 W, approx. 230–270 m.a.s.l.: Gómez et al., 2017). We broadly classified the habitat into inundated (floodplain) and non-inundated (terrace) forest. Mean annual rainfall is approximately 2770 mm and mean annual temperature is 23°C (Gómez et al., 2017). Weather data, including daily rainfall and daily mean temperatures, were available for the field station from the AABP Atrium (AABP Atrium, 2013). The study site is described in greater detail in our earlier paper (Gómez et al., 2017) and one of the datasets associated with the present paper (Hopkins, Gómez, et al., 2023).

Kibale National Park, in western Uganda, contains medium altitude moist evergreen forest as well as swamps, grasslands, woodland thickets and colonizing shrubs (Struhsaker, 1997; Chapman & Lambert, 2000), and is nowadays surrounded by agricultural land. Our study site was near the Makerere University Biological Field Station (0°33.750 N, 30°21.370 E; approx. 1500 m.a.s.l.). The area contains a varied mix of different habitats, which we broadly classified into a successional gradient from farmland and clearcut former plantation to primary forest (Hopkins, Roininen, & Sääksjärvi, 2019a; b). Mean annual rainfall is approximately 1700 mm, mean maximum daily temperature 24°C and mean minimum daily temperature 16°C (Chapman et al., 1999). Mean average temperatures are not available for the site, but are estimated to be 20°C by the CHELSA climate data set (Karger et al., 2017, 2021). Weather data consisting of daily rainfall and maximum and minimum temperatures were collected by a worker at the field station during the Malaise trapping. The study site is described in greater detail in our earlier paper and its associated dataset (Hopkins, Roininen, & Sääksjärvi, 2019a; b).

### Malaise trapping

We collected insects by Malaise trapping in the same way in both Peru and Uganda. Malaise traps were of a standard size and design: black with a white roof, approximately 170 cm long with two 1.6 m ^2^ openings, with identical fabric, mesh sizes and collecting jars, supplied by Marris House Nets (for Peru) or its successor B&S Entomological Services (for Uganda). The traps were placed on the likely flight paths of insects, and they collected flying insects into approximately 80% ethanol. In both Peru and Uganda, traps were used for a whole year to cover all seasons, and large numbers of traps were placed in different habitats. In Peru, three shorter sampling campaigns were also carried out. Traps were emptied at intervals mostly ranging from one to three weeks.

The Peruvian Malaise trapping consisted of a full year in 2000 (January 2000 – January 2001), and three shorter sampling campaigns in 1998 (August 1998 – January 1999), 2008 (May – August 2008) and 2011 (April – December 2011). In 1998, ten traps were placed in Allpahuayo-Mishana: four in clay soil forest and six in white sand forest. The total sampling effort was 45.8 trap months (c.f. 44 trap months mentioned in Sääksjärvi et al., 2004). In 2000, seventeen traps were placed in Allpahuayo-Mishana: two in clay soil forest, four in loamy soil forest and a total of eleven in three kinds of white sand forest (varying e.g. in canopy height and reflecting the high habitat heterogeneity of the study area). Fourteen of these traps were in place for the whole time period, but one trap (i3) was stolen in June 2000, and two traps (k1, k2) were installed as a replacement in July 2000. The total sampling effort was 151 trap months according to the compiled data (c.f. 141 trap months mentioned in Sääksjärvi et al., 2004). In 2008, nine traps were placed in Los Amigos: four in floodplain forest and five in terrace forest. The total sampling effort was 27.1 trap months. In 2011, fourteen traps were placed in secondary forest in Allpahuayo-Mishana. Only four of them were in place the whole time period: eight others were placed in October, one was stolen before it collected any samples, and one was stolen in August. The total sampling effort was 45.8 trap months (c.f. 45 trap months mentioned in Gómez et al., 2017). The full metadata of the traps has been published in the compiled dataset of Hopkins, Gómez, et al. (2023). This includes trap and sample data such as the time period when each sample was collected, the trap locations, and weather data. The 1998 and 2000 sampling campaigns have been described in greater detail by Sääksjärvi et al. (2004, 2006), and the 2008 and 2011 sampling campaigns by Gómez et al. (2017). In the present paper, we focus on the Rhyssinae data, which have not been published before.

The Ugandan Malaise trapping consisted of a full year, September 2014 – September 2015. A total of 34 traps were placed: sixteen in primary forest, seven in disturbed forest, nine in clearcut former plantations and two outside the natural park in agricultural land. The total sampling effort was 373.5 trap months, with a further 8.9 trap months being unrepresentative of a normal catch for various reasons, e.g. due to the traps and their samples being trampled by elephants. The Ugandan Malaise trapping data used in the present paper is from a previously published dataset (Hopkins, Roininen, & Sääksjärvi, 2019a). The sampling campaign has been described in greater detail by Hopkins, Roininen, & Sääksjärvi (2019b).

The original data from the Peruvian sampling campaigns had become partly fragmented over the years, so we recompiled the data from a variety of sources, such as old computer files and the labels on insect specimens and sample jars. The compiled dataset is available online (Hopkins, Gómez, et al., 2023). It includes a complete list of the Peruvian Malaise samples: what samples were collected, when they were collected and what trap they came from. The dataset also provides information on the trap sites (including vegetation near the traps) and on the weather during the Malaise trapping. The source material and files detailing how the data were compiled and inconsistencies resolved are also provided.

### Rhyssine wasps

The rhyssines and other ichneumonoid wasps (families Ichneumonidae and Braconidae) were separated from the Malaise samples, and are currently at the Zoological Museum of the University of Turku (ZMUT), Finland. All Peruvian wasps have been pinned and sorted to subfamilies. Ugandan wasps are still being processed, but all rhyssines have been pinned.

The Peruvian rhyssines had not been databased before this study. We databased a total of 94 individuals: 87 individuals found at the museum, and a further seven individuals that were mentioned in an earlier paper (Gómez et al., 2015). Four of these were left out of analyses due to it being unclear which sample they came from. We expect the effect of any rhyssines being missed during databasing to be small, since other potential sources of error (e.g. wasps lost during the processing of samples) are much greater. Other sources give the total as 96 rhyssines (Gómez et al., 2017) or 93 rhyssines (file 2 in folder “1 Raw Data” of Hopkins, Gómez, et al., 2023). Sääksjärvi et al. (2006, Appendix 1) reported 945 rhyssine+pimpline individuals caught in 2000, which is 8% more than in our data (874 rhyssines+pimplines in file 2 in folder “1 Raw Data” of Hopkins, Gómez, et al., 2023).

The rhyssine wasps were sorted into species at ZMUT. Peruvian rhyssines were identified to species mainly by Ilari Sääksjärvi and Isrrael Gómez, and the species delimitation was later verified by Tapani Hopkins. Ugandan rhyssines were identified to species mainly by Tapani Hopkins, and the species delimitation was verified by Ilari Sääksjärvi. Species delimitation was based on finding at least one morphological character (or combination of characters) unique to the species, backed up by differences in colouration. Colour was mostly not used as a morphological character, since in our experience it varies within wasp species. However, the colour of the hind wing was assumed to be species-specific for the Peruvian rhyssines, due to other distinguishing characters being unclear or otherwise hard to use for identifying specimens. The Peruvian species have been taxonomically reviewed by Gómez et al. (2015) and the Ugandan species by Hopkins, Roininen, van Noort et al. (2019).

To check if differences in how Peruvian and Ugandan species were delimited could have affected the results by inflating the number of Peruvian species, we created two additional Peruvian species delimitations. In the “semi-conservative” delimitation, *Epirhyssa zaphyma* Porter (Porter, 1978) and *E. lutea* Gómez & Sääksjärvi (Gómez et al., 2015) were treated as the same species. These species would likely not have been treated as different species if they had been caught in Uganda instead of Peru, since they are very similar and the two main characters that separate them (clypeus, tergite 1) varied greatly within Ugandan species. In the “conservative” species delimitation, the colour of the hind wing was also discounted as a character, and species were merged if they were not clearly separated by some additional character that was at least as clear as the characters used to separate Ugandan rhyssine species. This delimitation is overly conservative in Peru and merges obviously valid species, which makes it useful in giving an absolute lower bound to the Peruvian species count, irrespective of how species are delimited. In particular, it ignores subtle distinguishing characters (such as e.g. the proportions of wing vein lengths), many of which were too inconvenient for everyday species identification to be included in Gómez et al. (2015). In this conservative delimitation, the following species pairs were treated as if they were one species: *Epirhyssa zaphyma* and *E. lutea*; *E. diatropis* Porter (Porter, 1978) and *E. ignisalata* Gómez & Sääksjärvi (Gómez et al., 2015); *E. braconoides* Porter (Porter, 1978) and *E. cochabambae* Porter (Porter, 1978); and *E. pertenuis* Porter (Porter, 1978) and *E. iiapensis* Gómez & Sääksjärvi (Gómez et al., 2015) (this last pair turned out not to affect our results, as no *E. pertenuis* were found in our samples).

The data on all the rhyssine wasps used in the present paper is available in the supplementary dataset (Hopkins, Tuomisto, et al., 2023). This contains the full specimen data of all the rhyssines caught by Malaise trapping during the four Peruvian sampling campaigns and the Ugandan sampling campaign, and includes the place of deposition of each specimen.

### Analyses

We compared the Peruvian and Ugandan rhyssines using two counts: the number of individuals caught, and the number of species caught.

To examine how the number of rhyssine *individuals* caught in Peru and Uganda differed, we calculated the number of individuals caught during each sampling campaign per trap day (average number of individuals caught by one trap in one day) or trap month (30.5 trap days). This adjusts for the different sampling efforts.

Since rainfall often decreases Malaise trap catches, and it rained more in Peru than in Uganda, we estimated if observed differences in the number of individuals caught could be due to differences in rainfall. We estimated how many rhyssines would have been caught in Uganda if it had rained as much as it did during the main Peruvian sampling campaign in 2000. We did this by transferring Peruvian daily rainfall to the Ugandan data, then scaling down Ugandan catches based on how much the rainfall increased. We calculated the decrease in catches using the model in Hopkins, Roininen, & Sääksjärvi (2019b). This model is based on comparing the Ugandan rhyssine catches to weather data, and gives a separate estimate of the effect of rainfall for each species. This provides only a rough estimate, since we are extrapolating beyond the Ugandan data which the model is based on, but is sufficient for our purpose of checking if rainfall could plausibly be the cause of observed differences in Malaise sample abundances.

To compare the rate at which *species* were caught in Peru and Uganda, and thereby obtain an idea of overall species richness, we created species rarefaction curves. These show an estimate of how the number of observed species is expected to increase as a function of the number of individuals caught. Because the number of species observed is directly constrained by the number of individuals observed, and traps differed in how many individuals they captured per day, we used the observed number of individuals on the x axis of the rarefaction curves instead of the number of trap days (Gotelli & Colwell, 2011; Gómez et al., 2017). For resampling, each roughly two-week sampling interval was considered a sample, and these were resampled without replacement 100 times. In other words, we carried out a sample-based rarefaction (the individuals of each sample were kept together in resampling), but displayed the species accumulation results against the cumulative number of individuals rather than cumulative number of trap days. We produced separate species accumulation curves for each sampling campaign and forest type. For the Ugandan samples, we also produced separate species accumulation curves for the wet season and the dry season.

We compared four of the rarefaction curves in greater detail: the clay and loam soil curves of the main Peruvian sampling campaign in 2000, and the wet and dry season primary forest curves of the Ugandan sampling campaign. These sampling campaigns and forest types gave the largest sample sizes and were the most relevant to compare (e.g. the Peruvian white sand forest has no clear equivalent in Uganda). To roughly estimate whether the Peruvian and Ugandan curves differed significantly from each other, we calculated approximate 84% confidence intervals (these intervals overlap when p ≥ 0.05, see chapter 4.2.6 of Gotelli & Colwell, 2011). This should, however, be treated as a rough guide only, since we are calculating the intervals approximately (by resampling) instead of using unconditional variance as in Gotelli and Colwell (2011). Differences between rarefaction curves reflect differences in species diversity, which in turn consists of species richness and the evenness of species abundances (Gotelli & Colwell, 2011; Tuomisto, 2012). To find out to what degree the rarefaction results reflect differences in species richness (which is related to the size of the regional species pool) rather than in evenness, we quantified evenness ^*q*^*E* at *q*=1 and *q*=2. Evenness is calculated as ^*q*^*E*=^*q*^*D*/*R*, where *R* is species richness and ^*q*^*D* is diversity of order *q*. At q=1, ^*q*^D corresponds to Shannon diversity (exp(H’), where H’ is the Shannon diversity index) and at q=2 it corresponds to Simpson diversity (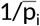, where 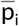 is the weighted arithmetic mean of the species’ proportional abundances; Tuomisto, 2012). The value of evenness ranges from 1/*R* to one, with higher values for more even abundances.

All analyses were carried out in the R software, v. 4.2.1 (R Core Team, 2017). The R code and data are available online (Hopkins, Tuomisto, et al., 2023).

## Results

Peruvian traps caught a total of 90 rhyssine individuals, which is only a fifth of the 444 individuals caught by Ugandan traps (Figures 1-2). The difference was partly due to a larger total sampling effort in Uganda (374 trap months) than in the four Peruvian sampling campaigns (a total of 270 trap months), but there was also a clear difference in the catch per unit time. The Peruvian traps caught 0.2, 0.4, 0.6 and 0.3 rhyssine individuals per trap month during the 1998, 2000, 2008 and 2011 sampling campaigns, respectively. None of these came even close to the average of 1.2 rhyssines per trap month caught in Uganda.

**Figure 1.**
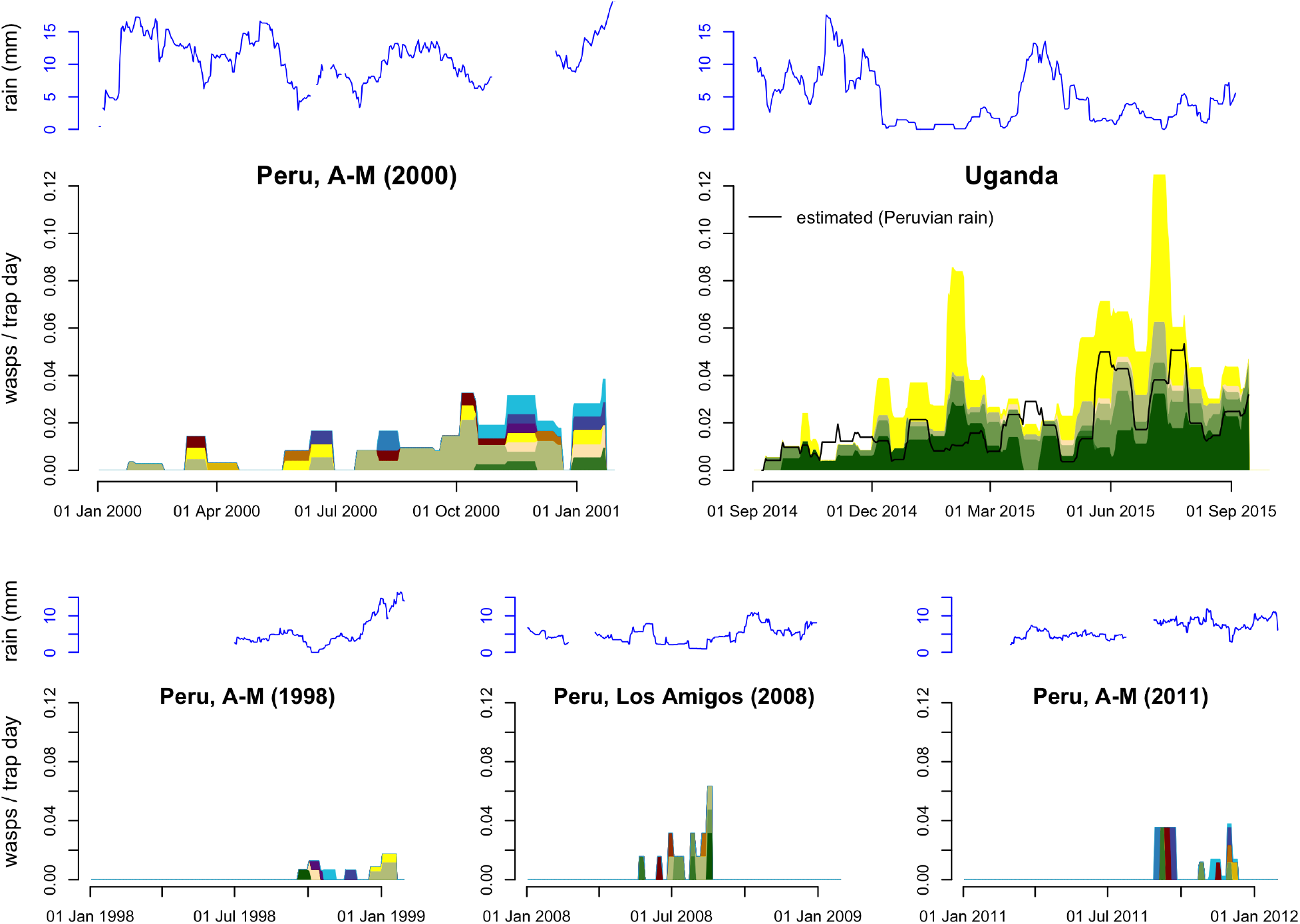
Rate of rhyssine captures in Malaise traps (individuals / trap day) operated in Peru and Uganda, with the amount of precipitation during the sampling period (29 day rainfall averages for Peru, 15 day averages for Uganda). The Peruvian sampling campaign 2000 in Allpahuayo-Mishana (A-M) provided most of the Peruvian data, the three shorter sampling campaigns provided additional data. Far fewer rhyssines were caught in Peru than in Uganda. However, when we estimated how many Ugandan rhyssines would have been caught if it had rained as much as during the Peruvian sampling campaign 2000 (black line), the difference was much smaller. The proportions of different species in the catches of each sample (which were accumulated over a time period of 1–3 weeks each) are visualised by colour differences (these are species-specific within but not among panels).

**Figure 2.**
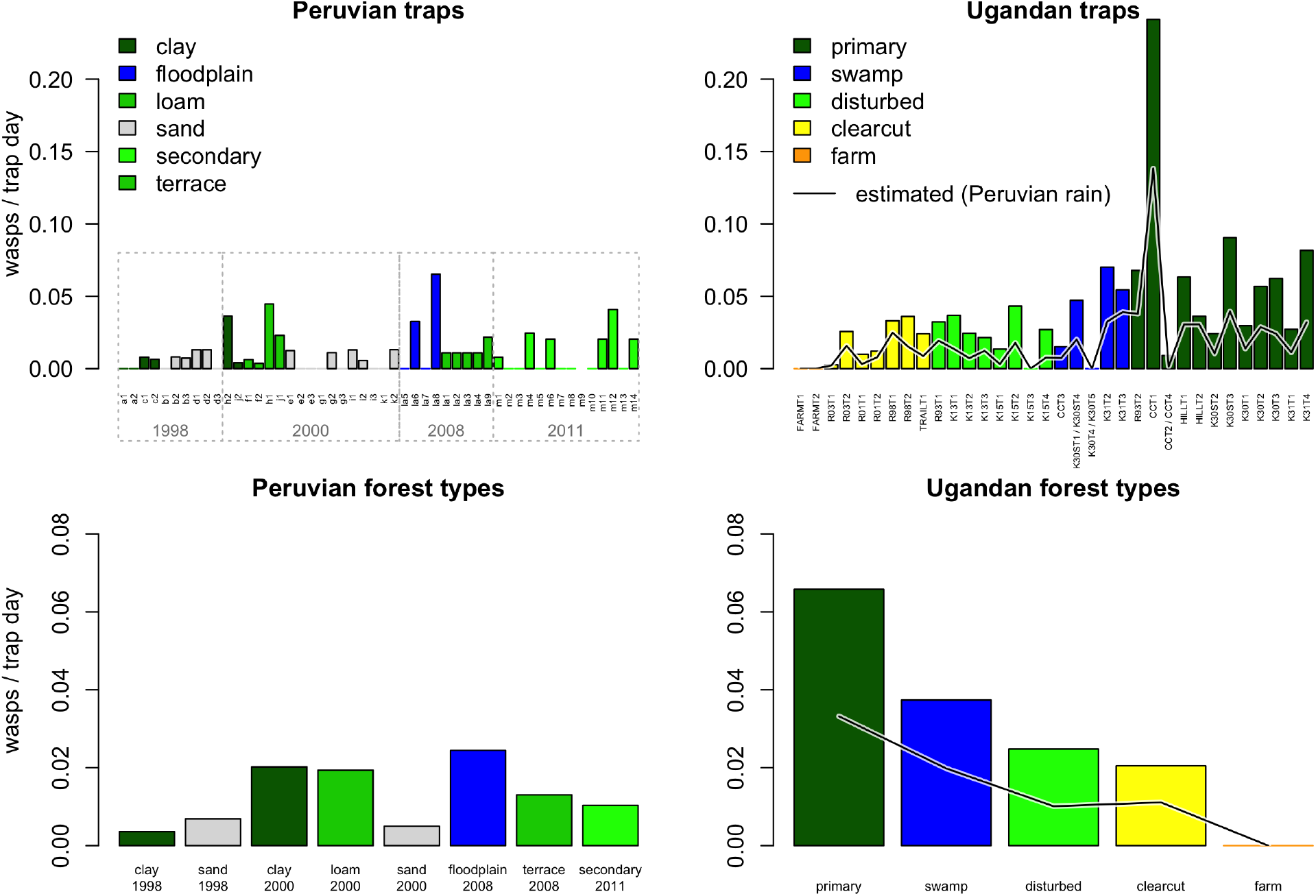
Number of rhyssines caught in 50 Peruvian and 34 Ugandan traps. The top panels show the catch (individuals per trap per day) for each trap, and the bottom panels the average catch for each forest type. Far fewer rhyssines were caught in Peru than in Uganda. However, when we estimated how many Ugandan rhyssines would have been caught if it had rained as much as during the Peruvian 2000 sampling campaign (black line), the difference was much smaller.

The difference between Peruvian and Ugandan rhyssine catch sizes was largely explainable by differences in rainfall (Figures 1-2). Using a regression model based on the Ugandan data, we estimate that a total of 220 Ugandan rhyssine individuals (0.6 per trap month) would have been caught if it had rained as much in Uganda as it did during the main Peruvian sampling campaign (2000). This is only half of what the Ugandan traps actually caught, but close to what the Peruvian traps caught.

The Peruvian Malaise trapping caught a total of 14 rhyssine species: 7, 11, 7 and 8 species for the 1998, 2000, 2008 and 2011 sampling campaigns, respectively. The Ugandan Malaise trapping caught only 6 species, despite a larger collecting effort in terms of trap months and a much larger number of individuals caught. The difference was not caused by differences in how species were delimited: the Peruvian species counts for the four sampling campaigns were 7, 10, 7 and 7 species for a semi-conservative species delimitation (with a total of 13 species), and 6, 9, 6 and 6 species for an extremely conservative species delimitation (with a total of 11 species).

The Peruvian Malaise trapping accumulated rhyssine species faster (per individual caught) than did Ugandan Malaise trapping, irrespective of forest type (Figure 3). Although there was some variation in how quickly species accumulated, Peruvian and Ugandan rarefaction curves clearly fell outside each other’s confidence intervals once a sufficient number of individuals were included (Figure 4). However, Peruvian sample sizes were relatively small (Figures 4–5) and most Peruvian traps only caught a few individuals. The relative abundances of the species followed a similar distribution in Peruvian and Ugandan forest types, with the Peruvian species of some forest types less evenly distributed (Figure 5). Different forest types generally contained the same rhyssine species in Uganda, whereas there was more differentiation between forest types in Peru (Figure 5). No Peruvian trap caught more than seven of the 11–14 species, and no Ugandan trap more than five of the six species.

**Figure 3.**
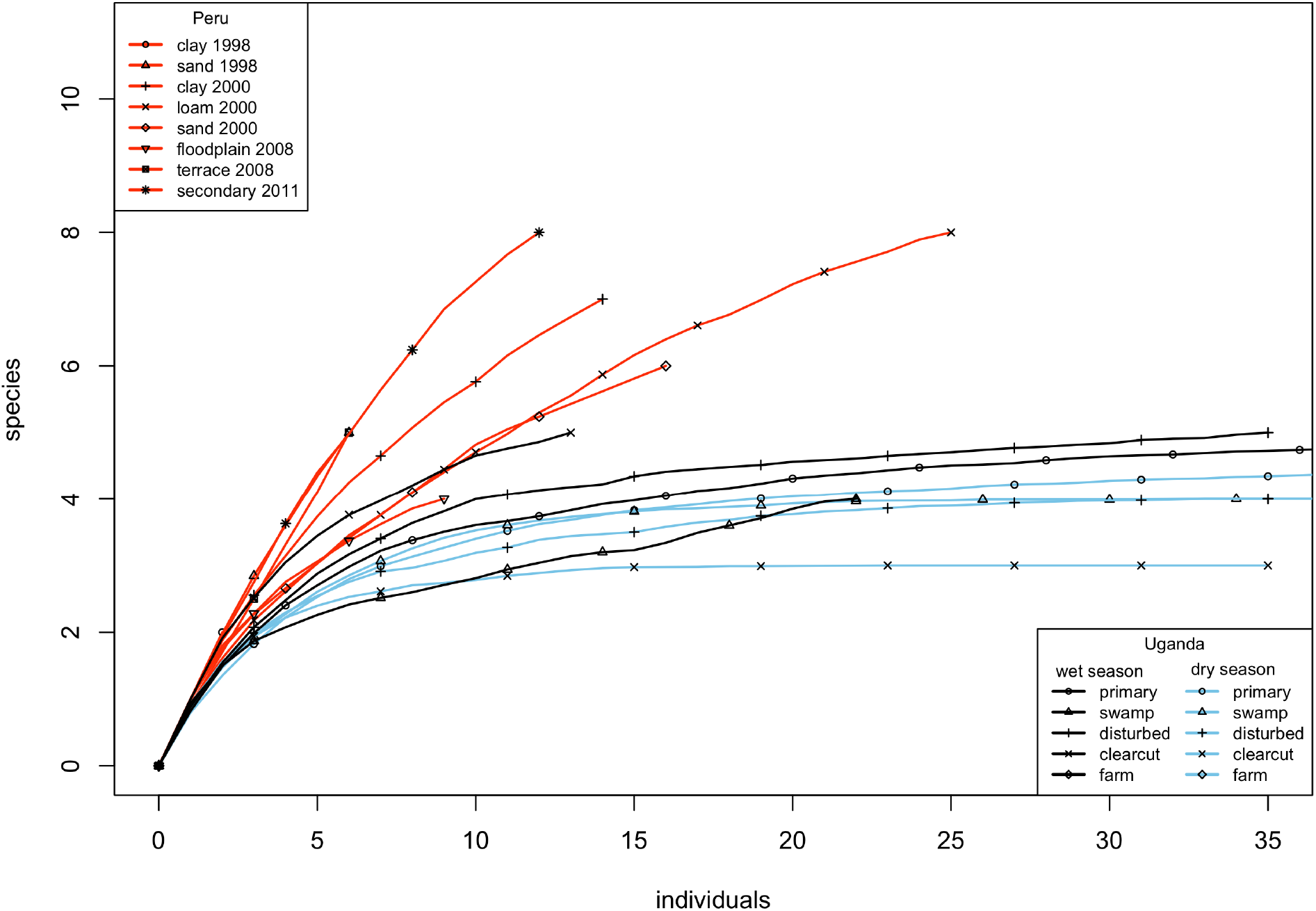
Sample-based species rarefaction curves showing how quickly Malaise traps caught species in Peru and Uganda. Each sample consists of a sample jar representing a sampling interval of mostly between 1 and 3 weeks. Peruvian curves were constructed separately for each forest type and sampling campaign (2008 sampling was at the Los Amigos site, others at Allpahuayo-Mishana). Ugandan curves were constructed separately for each forest type and season.

**Figure 4.**
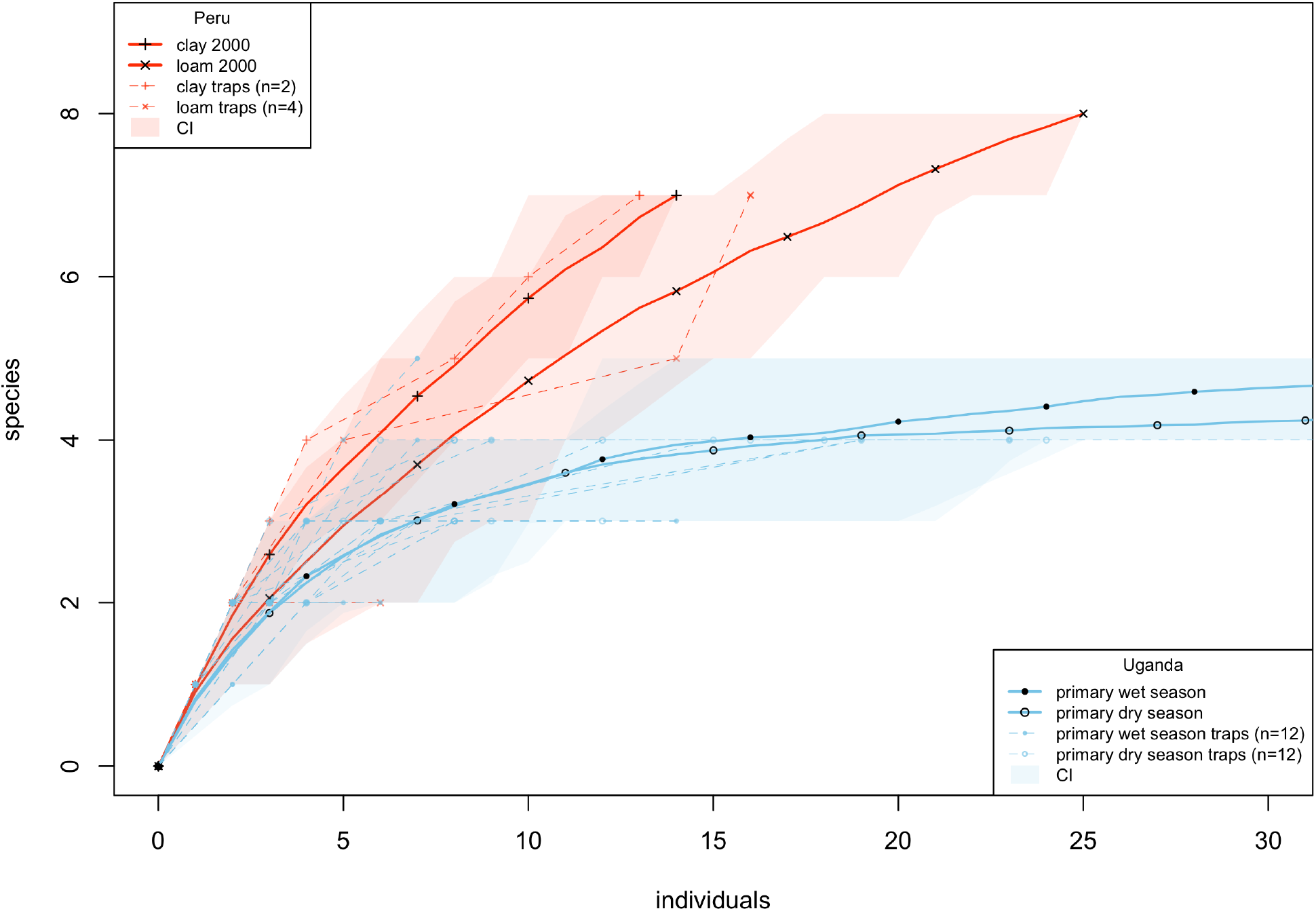
Selected species rarefaction curves showing how quickly Malaise traps caught species in Ugandan non-inundated primary forest, and in the closest equivalent to this forest type in Peru (main sampling campaign 2000). Shaded areas are confidence intervals that show how much variation there was in a curve (84% of a curve’s resamples fell inside its shaded area). The shaded areas of Peruvian and Ugandan curves do not overlap after 16 individuals, which suggests that the difference between them is significant. The accumulation curves of individual traps are also shown: Peruvian sample sizes were relatively low, with only two of the Peruvian traps in this figure catching more than six individuals.

**Figure 5.**
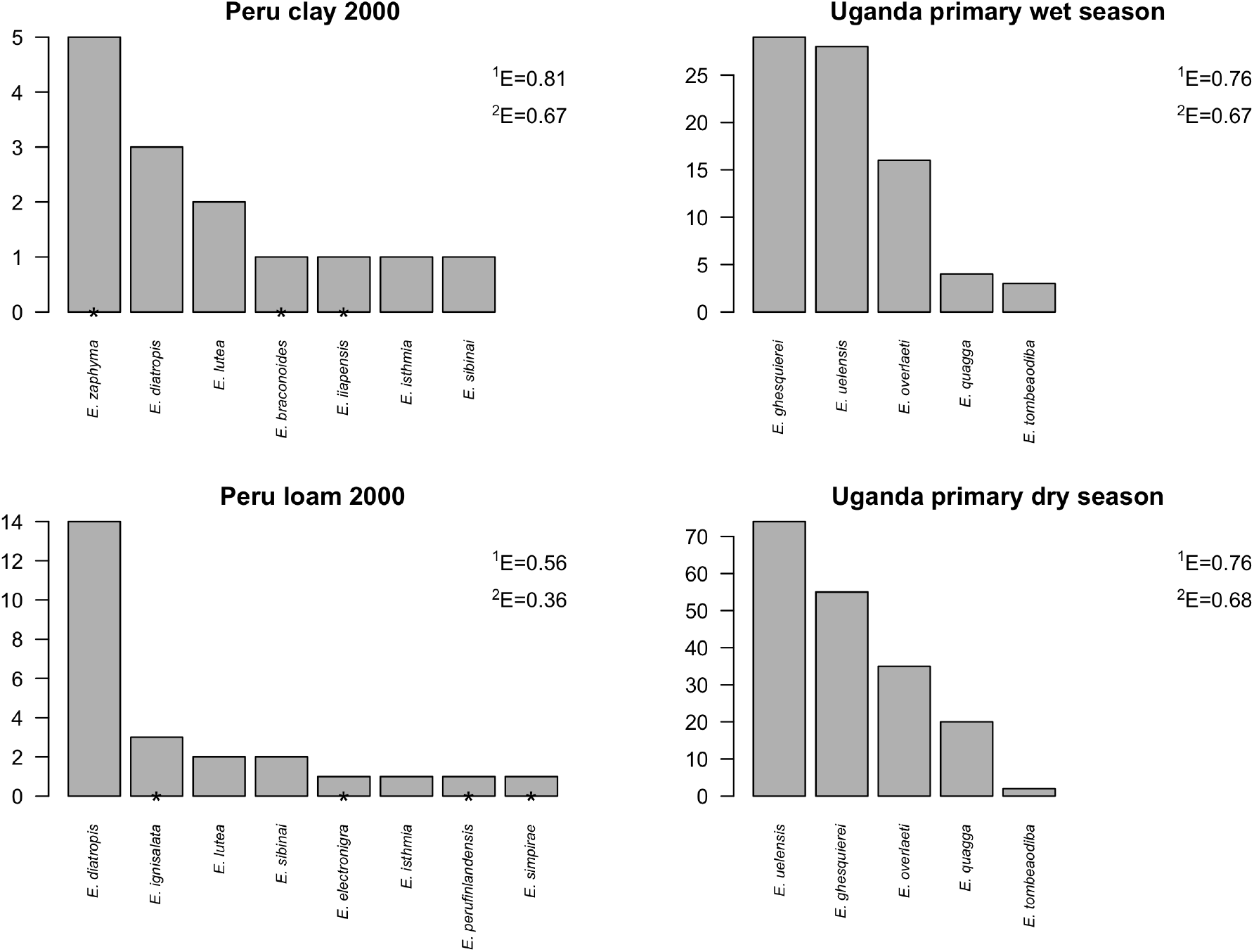
Number of individuals of each species caught in non-inundated primary forest in Uganda, and in the closest equivalents to this forest type in Peru (main sampling campaign 2000). Species that were only found in one of the forest types are marked with ^*^. The species abundance distributions were approximately equally even in the Peruvian clay soil sites and the Ugandan sites, but the Peruvian loam soil sites showed a smaller evenness value, indicating a higher degree of dominance (evenness values shown to the right of each panel).

## Discussion

### More species but fewer individuals in Peru

We observed two main differences between the Peruvian and Ugandan Malaise trapping results: Ugandan traps caught many more rhyssine individuals per unit time, but Peruvian traps caught more species. The observed difference in the number of rhyssine individuals caught reflects differences in rainfall. Peru had more rain than Uganda, and rainfall has been observed to decrease the number of rhyssines caught by Malaise traps, mostly by decreasing the rhyssines’ flight activity (although there are signs that abundances may also be affected: Hopkins, Roininen, & Sääksjärvi, 2019b). We estimated that if it had rained as much in Uganda as in Peru, Ugandan catches would have been halved and would have been much more similar to the Peruvian catches (Figures 1-2). The limitations of this estimate must be born in mind, however: it is a rough estimate based on extrapolating beyond our Ugandan data. Nevertheless, the role of rain is supported by two other observations. First, other taxa also seem to have been affected. Peruvian Malaise samples generally seem to be about a half or one third of the volume of Ugandan samples (based on observing how full the sample containers are after similar lengths of trap operation), which suggests that they contain fewer insects overall. Second, even within Peru, the drier site Los Amigos captured more individuals per trap month than the wetter Allpahuayo-Mishana did (0.6 versus 0.2–0.4 individuals per trap month, respectively). It thus seems plausible that rainfall, rather than some other ecological factor, is the main reason why Amazonian traps catch fewer individuals than Ugandan traps.

We speculate that there could be interesting ecological consequences of rain decreasing the flight activity, and possibly also abundance, of rhyssines: this could favour wood-boring insect larvae (which are the likely hosts of tropical rhyssines) in rainy areas as compared to drier ones. If Malaise traps encounter fewer rhyssines due to rain, wood-boring larvae probably do so too, and thereby face less predation pressure. Although there are other taxa that potentially compete with rhyssines for hosts, and could compensate for the lower predation pressure, many of them are likely also affected by rain. These include e.g. *Apechoneura* (Labeninae), *Dolichomitus* (Pimplinae), *Anastelgis* (Pimplinae) and woodpeckers (Picidae). This hypothesis could be tested by measuring the relative densities of infected and uninfected wood-boring larvae in Peruvian and Ugandan decaying wood. For our study sites, we would expect the highest densities of uninfected wood-boring larvae in Allpahuayo-Mishana, followed by the somewhat drier Los Amigos, and much lower densities at our Ugandan site. It is likely that other factors (such as humidity of the wood) also affect the densities of wood-boring larvae, so further studies would be needed to test this hypothesis.

Our results strongly suggest that Allpahuayo-Mishana in Peruvian Amazonia has more rhyssine species than Kibale National Park in Uganda. The species accumulation curves show that by the time 15 individuals had been caught, the mean observed number of species in the Ugandan traps had near-stabilised at little more than 4, whereas in the Peruvian traps the number continued to grow although it had already exceeded 6. The observed number of species reflects species diversity, which in turn depends on both the actual number of species present in the community (richness) and how similar the proportional abundances of the species are (evenness). In highly uneven communities, more individuals need to be sampled before the rare species get detected. The relative abundances of different species in our samples (Figure 5) suggest that evenness does not explain the differences we observed. Species accumulated slower in the Ugandan traps even though evenness in them was roughly the same as in the Peruvian clay soil traps, and greater than in the Peruvian loam soil traps.

Our second Peruvian site, Los Amigos, may also have more species than Kibale in Uganda. However, the evidence is insufficient since only 15 rhyssine individuals were caught in Los Amigos. It is noticeable that the species did not accumulate as quickly in Los Amigos as in Allpahuayo-Mishana (Figure 3), which could indicate that the total species richness in Los Amigos may be somewhat smaller.

One potential factor affecting the number of species observed at different sites is the geography of the sites and how that geography has been covered. We sampled a very restricted area in Allpahuayo-Mishana (approx. 4 km × 4 km in year 2000, in other years no larger), Los-Amigos (approx 3 km × 3 km), and Kibale, Uganda (approx 4 km × 7 km). This will undoubtedly have restricted the pool of species we were able to discover. In Kibale, for example, the vegetation is known to vary greatly along a north-south gradient (Chapman & Lambert, 2000). Had we placed our traps nine kilometres further north at Sebitoli, or seven kilometres further south near Ngogo, they would have been in noticeably different forest, with even the dominant tree species partly differing, and potentially with different rhyssine species. More importantly, we also know that within our sites, not all habitats were sampled. Allpahuayo-Mishana in particular is known to have a large number of geologically and floristically distinct habitats in a relatively small area (Whitney & Alonso, 1998; Sääksjärvi et al., 2004), and we know of habitats such as papyrus swamp in Kibale which were not covered by our Malaise trapping. To what extent this affects the number of species caught depends on how strongly rhyssines are restricted to specific habitats. It is noticeable that in our Peruvian Malaise trapping, traps in different forest types caught partly different species (e.g. Figure 5), whereas our Ugandan traps largely sampled the same set of species. This may, however, be caused by the relatively small number of rhyssines caught in Peru. Overall, the relatively high diversity of different habitats at Allpahuayo-Mishana could be one factor explaining the larger number of species observed there.

If the apparent greater rhyssine species richness of Allpahuayo-Mishana is genuine, and reflects the situation of the rest of lowland Peru, it would match what little is known for other taxa. In general, Neotropical forests are believed to be have more species rich floras than Afrotropical forests (Gentry, 1982), and the global species richness of many taxa seems to peak in western Amazonia: reptiles in Allpahuayo-Mishana (Gentry, 1988; reptiles listed in Dixon & Soini, 1975, 1977), trees in A-M (Vásquez Martínez & Phillips, 2000), birds (Pearson, 1977), butterflies in Tambopata, near Los Amigos (Gentry, 1988; butterflies listed in Lamas, 1984). It should also be noted that our Ugandan site is at a higher altitude than our Peruvian site, and species richness is generally thought to decrease with increasing altitude as well as latitude (Wolda, 1987; Fernandes & Price, 1988); although some taxa may peak at mid altitudes instead, at least where mid-altitudes have higher humidity and rainfall (Brehm et al., 2007; but see Molina-Martínez et al., 2013).

### Future Malaise trapping

Long-term, standardised Malaise trapping showed promise for global comparisons of species richness. Our current results involve only one (relatively rare) subfamily and just a few sites. However, the fact that we could get reasonable results demonstrates the potential of our method for drawing more far-reaching conclusions. Sampling more sites in both tropical Africa and Amazonia, for example, and analysing all (or at least the most abundant) subfamilies, would give an estimate of which continent has more parasitoid wasp species. It would also detect differences in the relative abundance and species richness of different subfamilies. Such sampling could especially focus on low altitude equatorial sites (e.g. the Democratic Republic of the Congo in Africa) or include a variety of altitudes on every continent.

Although our sampling design with standardised Malaise trapping succeeded in getting comparable data, further improvements could be made for future Malaise trapping. These include greater focus on covering all habitats at a site and using modelling approaches to generate easy-to-compare data.

Our trap placement was unbalanced between habitat types: if we had placed more Peruvian traps in loam or clay forest, for example, we would have obtained a better coverage of what appear to be highly variable habitats. These habitats were also the ones with a clearest equivalent in Uganda. As it was, we only had six such traps in our main Peruvian sampling campaign. This does not give a good coverage of the kind of varied habitat that could well be subdivided into more than six habitat types (how habitat is classified into habitat types is to some extent a matter of preference). Ideally, since Malaise traps typically give very variable catches even in the same habitat (Fraser et al., 2008; Saunders & Ward, 2018; Chimeno et al. 2023), every habitat type should be covered by several traps. In practice, this requires too many traps to be feasible. We do not have a full solution to this problem, but suggest that future inventories devote effort to obtaining as good a habitat classification as possible at an early stage. Teaming up with a botanist when planning where to place traps, for example, would allow the available traps to be placed optimally (see e.g. Sääksjärvi et al., 2006). Many plant taxa such as ferns have been found to be good indicators of tropical forest habitat types (Salovaara et al., 2004; Pomara et al., 2012; Zuquim et al., 2014). Even the simple expedient of photographing ferns at trap sites, then showing the photographs to a specialist, could help classify the sites into habitat types (Suominen et al., 2015).

Having enough traps in each habitat, together with getting a sufficient sample size and coverage of all seasons, allows modelling the expected catches of traps (i.e. how many individuals to expect for a given habitat and weather). This could potentially be a great advantage when comparing two different sites: without a model, we are comparing Malaise samples which stem from a varying mix of different habitats, have been collected during varying weather, and are otherwise hard to treat statistically. In this work, we used a model of how Ugandan rhyssines react to rainfall to account for the effect of different rainfall in Peru and Uganda. Unfortunately, the Peruvian sample sizes (total of 90 rhyssine individuals, split among 14 species) were too low to allow modelling of the Peruvian rhyssines. Other subfamilies have mostly not yet been fully processed (except for Peruvian Pimplinae: Sääksjärvi et al., 2004; Gómez et al., 2014), but many should have much larger sample sizes than Rhyssinae and would be modellable. Wherever possible, we suggest building a model of how many individuals to expect in a given habitat and weather, then comparing the models of different sites instead of the raw data (Malaise samples) from which the models have been interpolated.

## Acknowledgements

Heikki Roininen supervised and otherwise supported the Ugandan Malaise trapping. Isaiah Mwesige helped maintain the Ugandan Malaise traps and ably carried out a whole range of other fieldwork. Yosinta Tumusiime gathered the Ugandan weather data. Our field research was supported by the staff of the Makerere University Biological Field Station. Countless people helped process both the Peruvian and Ugandan samples, including the staff of the Zoological Museum of the University of Turku, students of the university and school pupils from throughout the Turku region and students of the Universidad Nacional de la Amazonía Peruana (UNAP, Peru). These contributions are gratefully acknowledged.

The required research and export permits were issued by the Uganda National Council of Science and Technology (NS 504) and the Uganda Wildlife Authority. The Ministry of Agriculture and the Ministry of Environment of Peru provided the collecting and export permits for the Peruvian samples. We appreciate the support of the Instituto de Investigaciones de la Amazonía Peruana (IIAP), Universidad Nacional de la Amazonía Peruana (UNAP), the Amazon Conservation Association (Peru), Conservation International Foundation (Peru) and the late Pekka Soini for the field studies made in Allpahuayo-Mishana and Los Amigos.

## Data, scripts, code, and supplementary information availability

The analyses and data are available online in two datasets. The first of these contains the code and data used in the statistical analyses: https://doi.org/10.5281/zenodo.8208661 (Hopkins, Tuomisto, et al., 2023). The second contains the data on the Peruvian Malaise trapping, and a description of how it was compiled: https://doi.org/10.5281/zenodo.8208660 (Hopkins, Gómez, et al., 2023).

